# Manufacturing Stable Bacteriophage Powders Using Thin Film Freeze-drying Technology

**DOI:** 10.1101/2020.11.27.401505

**Authors:** Yajie Zhang, Melissa Soto, Debadyuti Ghosh, Robert O. Williams

## Abstract

Recently, therapeutic uses of bacteriophage (phage) are gaining increased attention, yet common liquid phage formulations require cold chain storage that limits their potential use. Phage therapy is considered as an alternative to antibiotics for bacterial infections and more significantly a promising solution for the ever-increasing prevalence of multi-drug resistance (MDR) pathogens. One of the most promising applications of this therapy is to treat pulmonary bacterial infections. To efficiently deliver therapeutic phage to the lungs, phage formulations that allow for nebulization or dry powder inhalation are under active development. Several conventional particle engineering technologies have been applied in the development of dry powder inhalers (DPI), including spray drying, spray freeze drying, and atmospheric spray freeze drying, but these processes have their own disadvantages that limit their use with bacteriophage formulations and delivery. In our work, we hypothesize that thin film freeze-drying (TFFD) can be used to produce brittle matrix powders containing phage that may be suitable for delivery by several routes of administration, including by nebulization after reconstitution and by intranasal or inhalation delivery of the resulting dry powder. Here we selected T7 bacteriophage as our model phage in a preliminary screening study and found that a binary excipient matrix of sucrose and leucine at ratios of 80:20 or 75:25 by weight, protected bacteriophage from the stresses encountered during the TFFD process. In addition, we confirm that incorporating a buffer system during the TFFD process significantly improved the survival of phage during the ultra-rapid freezing step of the TFFD process and subsequent sublimation step in the lyophilization process. This preservation of phage bioactivity was significantly better than that observed for formulations without a buffer system. The titer loss of phage in standard SM buffer (Tris/NaCl/MgSO_4_/gelatin) containing formulation was as low as 0.2 log plaque forming units (pfu), which indicates that phage functionality was preserved after the TFFD process. Moreover, the presence of buffers markedly reduced the geometric particle sizes as determined by a dry dispersion method using laser diffraction, which indicates that the TFFD phage powder formulations were easily sheared into smaller powder aggregates, an ideal property for facilitating pulmonary delivery through DPIs. From these findings, we show that TFFD is a particle engineering method that can successfully produce phage containing powders that possess the desired properties for bioactivity and inhalation therapy.

## Introduction

The rise of multidrug resistant bacteria and the resulting decline in effective small molecule antibiotics to treat severe bacterial infections remains a public health threat [1]. The urgency of finding an alternative therapeutic strategy has brought the previously overlooked bacteriophage (or phage) therapy back to researchers’ attention [2-4]. Phages are viruses that infect and suppress and/or eradicate specific strains of pathogenic host bacteria [5]. The advantage of phage therapy compared to antibiotics includes but is not limited to its 1) specificity to the target bacterial strain, allowing for minimal disturbance to normal human flora; 2) host bacterial based replication: the delivered phage relies on the presence of target bacteria in order to replicate 3) minor in vivo toxicity; 4) biofilm clearance ability; and importantly, 5) adaption to drug resistant bacteria [6].

However, to successfully develop phage into a pharmaceutical product, the shelf-life of the formulation should be prioritized. Since phage are mainly assembled with protein capsids which encapsulate genetic material, phage are commonly formulated with protein formulation strategies. In general, dry state formulations are considered more stable than their liquid counterparts [7-9]. In an aqueous environment, proteins can be stressed by different factors, such as mechanical agitation, pH shifts, thermal stress (induced by both low and high temperatures), surface tension, hydrolysis and deamination [10]. Solid state formulations can remain stable by avoiding the issues listed above. Therefore, formulating phage in a dry powder form can enhance its druggability by improving its storage stability and reducing its handling restrictions needed to retain its bioactivity.

Currently, phage formulations are mainly limited to liquid suspensions, which usually requires a cold supply chain. Excitingly, however, numerous studies have proven that phage are amenable to solid state formulations. Spray drying and lyophilization are two commonly used technologies reported to dehydrate phage formulations. For example, Alfadhel *et al*. formulated a strain of Siphoviridae phage with hydroxypropyl methylcellulose (HPMC) with and without the addition of mannitol, and the formulation was subsequently lyophilized into a dry powder [11], but HPMC is not used in inhaled dosage forms. Merabishvili *et al*. investigated the influence of different excipients on the stability of shelf freeze-dried phage (e.g., slow freezing rate on the conventional lyophilizer shelf). Interestingly, their data demonstrated that 0.5 M trehalose was the best formulation in terms of phage titer loss, which was 1.5 log at a 4 °C storage temperature after more than three years [12]. No studies were reported at storage temperatures greater than 4 °C. Matinkhoo et al. reported on spray dried Φ KZ/D3 phage powders and showed that less than a 0.15 log titer loss was found in all formulations after three months of refrigerated storage [13]. Leung et al. have also demonstrated that a spray dried powder matrix containing no less than 40% trehalose achieved suitable aerosol performance and preserved *Pseudomonas* phage activity [14]. Similarly, Chang et al. and others have reported on the development of inhalable phage dry powders by spray drying [15-19]. In addition to conventional lyophilization and spray drying, several alternative methods have been explored, including spray freeze-drying and atmospheric-spray freeze drying [20, 21].

Phage formulations in the dry state offer more options for delivery routes and dosage forms with higher patient compliance, such as dry powder inhalation (DPI), oral dosage forms, topical pastes and dusting powders [22-24]. For instance, phage formulations for pulmonary delivery to treat lung infections have been previously reported, particularly for chronic *S. aureus* and *P. aeruginosa* infections associated with cystic fibrosis (CF) [16, 20, 25]. Dry powder inhalation products offer better patient compliance compared to nebulizers because they are smaller, portable, do not require electricity for operation or require regular disinfection, demands less time for administration and do not require the patient to coordinate between breathing and actuation of the device [26]. Yet, to use DPIs for phage administration, stable phage powders with desirable aerosol performance are needed. Chang *et al*. employed Taguchi experimental design with funneling approach and demonstrated that lactose provided the best phage stability among trehalose, lactose, and leucine and that these spray dried formulations produced over 50% fine particle fraction desired for inhalation [18]. However, risks associated with the use of spray drying and accompanying solvent removal include protein unfolding, reversible or irreversible aggregation, and chemical degradation. These risks can ultimately limit the use of spray drying to prepare phage powder formulations. Also, the spray nozzle used in the spray drying process to atomize the liquid into small droplets with a high surface area to volume ratio, may expose the protein solution to mechanically induced stress which can ultimately affect protein stability [27]. Golshahi *et al*. reported that endotoxin-removed bacteriophages KS4-M and FKZ lyophilized with a 60:40 (w/w) lactose:lactoferrin matrix were successfully aerosolized and viable upon delivery to the lungs [28]. While this is an example of phage retaining viability post-lyophilization, protein stability can still be affected by the slow rate of freezing encountered using conventional shelf lyophilization, limiting its use in preparing phage powder formulations. Moreover, to make the solid dry powder suitable for inhalation, the lyophilized powder was milled after lyophilization, which exposed the phage to harsh stresses after being subjected to stresses during slow freezing and sublimation steps.

Thin film freeze drying (TFFD) utilizes ultra-rapid freezing, resulting in the formation of brittle matrix powders containing the active therapeutic. TFFD technology employs low temperatures ranging between -70°C and -180°C without imparting high shear stress on the drug formulation, thereby preserving heat/mechanical-sensitive therapeutic moieties, such as phage. Spray drying, which uses spray nozzles to atomize liquid drug formulations for conversion to dry powders, requires high temperatures to evaporate the formulation solvents. In contrast, TFFD is operated at ultra-low temperatures, uses rapid freezing rates, and does not require mechanical shearing. Similarly, SFD and ASFD also require high shear spray nozzles in order to atomize formulations into small droplets (creating a high surface area to volume ratio of the droplets) of the liquid into air above the surface of liquid nitrogen, after which the partially frozen droplets impact the surface of the liquid nitrogen. Exposure to these SFD and ASFD processing conditions can be detrimental to formulating proteins and phage. In contrast, TFFD freezes liquid formulations by creating a thin film on a cryogenic surface, eliminating the high shear air-liquid interface as well as allowing for more homogenous freezing. In this study, we hypothesize that TFFD can be used to generate brittle matrix powders containing phage that are physically and chemically stable, thus preserving phage activity.

## Materials and Methods

### 1. Phage amplification

Wild type T7 phage (T7Select^®^) and host BL21 *Escherichia coli* bacteria strain were purchased from Millipore Sigma (Burlington, MA, US). To prepare a sufficient concentration of phage for experiments, phage were amplified according to manufacturer’s protocol. Briefly, T7 phage were added to BL21 *E. coli* liquid cultures (OD600 of 0.2-0.3) at a multiplicity of infection (MOI) of 0.001-0.01 and amplified for 1-3 hours at 37 °C at 250 rotations per minute (rpm) (MaxQ4000 shaker, Thermo Fisher Scientific) until lysis was observed. Bacterial lysate was collected, clarified with 5M NaCl/LB and spun down at 10,000 rpm in a Sorvall XFR Centrifuge (Thermo Fisher Scientific, Waltham, MA, US) for 30 minutes at 4 °C. The supernatant containing the phage was collected, and phage were further precipitated by incubating phage samples with a 50% PEG 8000 solution overnight at 4 °C. Once precipitated, the phage was pelleted by spinning down at 14,000 rpm and resuspended in either phosphate saline buffer (PBS) or standard SM buffer and collected in 1.5 mL microcentrifuge tubes. To further purify the phage, a second PEG precipitation step was performed with the resuspended phage by precipitating with 50% PEG 8000 solution on ice for at least 30 minutes. This lysate-PEG mixture was then centrifuged at 14,000 rpm for 30 minutes and the resulting phage pellet was resuspended in 50-100 microliters (μL) of either PBS or SM buffer. Amplified phage were quantified by standard double-layer plaque assay and stored at 4°C.

PBS (pH 7.2-7.4) was purchased from Sigma-Aldrich (St. Louis, MO, US), and SM buffer was prepared according to the SM buffer recipe provided by Cold Spring Harbor Protocols. For SM buffer preparation, NaCl (Thermo Fisher Scientific, Waltham, MA, US), MgSO_4_·7H_2_O (Thermo Fisher Scientific, Waltham, MA, US) and Tris-HCl were combined at final concentrations of 100 mM, 8 mM, and 50 mM, respectively. A final pH of 7.4-7.6 was achieved by adjusting the ratio of Trizma^®^ and Tris-HCl (Sigma-Aldrich, St. Louis, MO, US). Gelatin was not added in this SM buffer.

### 2. Phage viability assay

The lytic bioactivity of phage, representing the amount of viable phage in both solution and powder samples, was assayed by performing a standard double-layer plaque titering assay. Phage samples were prepared in 10-fold serial dilutions using LB media. Ten μL of each dilution were then added to 200 μL of BL21 bacteria (OD600 = 1.0) followed by adding 1 mL of melted LB top agar. After this solution was briefly vortexed, the mixture was plated onto pre-warmed 6-well plates previously prepared with 5 mL of LB agar. Plates were incubated at 37°C for approximately 3-4 hours or at room temperature overnight, until plaques were visible for counting and quantification. Dilutions that resulted in between 30-200 plaques/well were considered reliable and counted. Titer loss was calculated by dividing the initial input phage concentration (i.e. titer) for each formulation solution by the resulting phage concentration of each sample. The reliability of this titering method was evaluated by performing one-day repeatability and day-to-day repeatability analyses (N=3 respectively). There was no statistical difference in the titering results between these two studies. The relative standard deviation (RSDs) of those two repeatability studies and the combined RSD (N=6) were all less than 0.02%.

The phage powders were reconstituted in sterile water to a final concentration of 10 mg/mL. For the viability test with films collected during the freezing step, the frozen thin films were thawed at room temperature in a capped vial before tittering.

### 3. Formulation preparation

Several excipients that are commonly used in solid phage formulation research were selected, including three disaccharides (lactose, sucrose, and trehalose), and one amino acid (leucine). The sugars were incorporated in the formulations either alone or combined with leucine to form a binary excipient matrix. The sugar: leucine ratios were 90:10, 75:25, and 60:40. The formulation solutions were prepared to have a solid content of 0.5% (w/v) which corresponds to solution concentrations of 5 mg/mL. Solid content refers to the weight to volume concentration of all components in the pre-process solution formulation. The titers of amplified phage stocks used for these studies were approximately 5 x 10^11^ plaque forming units (pfu)/mL. These stocks were added to formulations and diluted 1,000-fold to achieve final titers of approximately 5 x 10^8^ pfu/mL. Formulations were prepared in either PBS, SM buffer, or water.

### 4. Manufacturing phage powder by thin film freeze drying (TFFD)

The setup of the thin film freezing apparatus is shown graphically in Diagram 1. Each phage solution was passed through a standard 5 mL or 10 mL syringe at a rate of 30-50 droplets per minute. The droplets fell from a height of 10-15 cm above an absolute-flat bottom stainless-steel container that was pre-chilled by submerging it in liquid nitrogen. As a result of thermal conductivity through the steel, the resulting equilibrium surface temperatures were below the freezing points of the solutions and could reach temperatures as low as -100 °C. The working temperature was controlled by adjusting the height of the container in the liquid nitrogen and maintained between -65 to -75 °C, which was monitored with a thermocouple. Upon touching the surface of the stainless-steel container, droplets deformed into thin films and froze instantaneously. The frozen thin films were removed from the surface by a stainless-steel blade and further collected by submerging the thin films in liquid nitrogen. For each formulation, the films and liquid nitrogen were poured into a lyophilization vial, which was then covered to prevent particles from exiting the vial during the sublimation step of freeze drying. Finally, the vials were transferred directly to a −80°C freezer to store vials until being placed into the lyophilizer and to allow for any excess liquid nitrogen to evaporate.

A Virtis Advantage Lyophilizer (The Virtis Company, Inc., Gardiner, NY) was used to dry the frozen thin films. Primary drying was carried out at −40°C for 2,000 min at 100 mTorr and secondary drying was carried out at 25°C for 1,250 min at 100 mTorr. A 12 h linear ramp of the shelf temperature from −40°C to +25°C was used at 100 mTorr between these two drying steps. After the cycle was done, the containers were capped tightly and then stored in a vacuum chamber immediately after being removed from the lyophilizer.

**Diagram 1.**
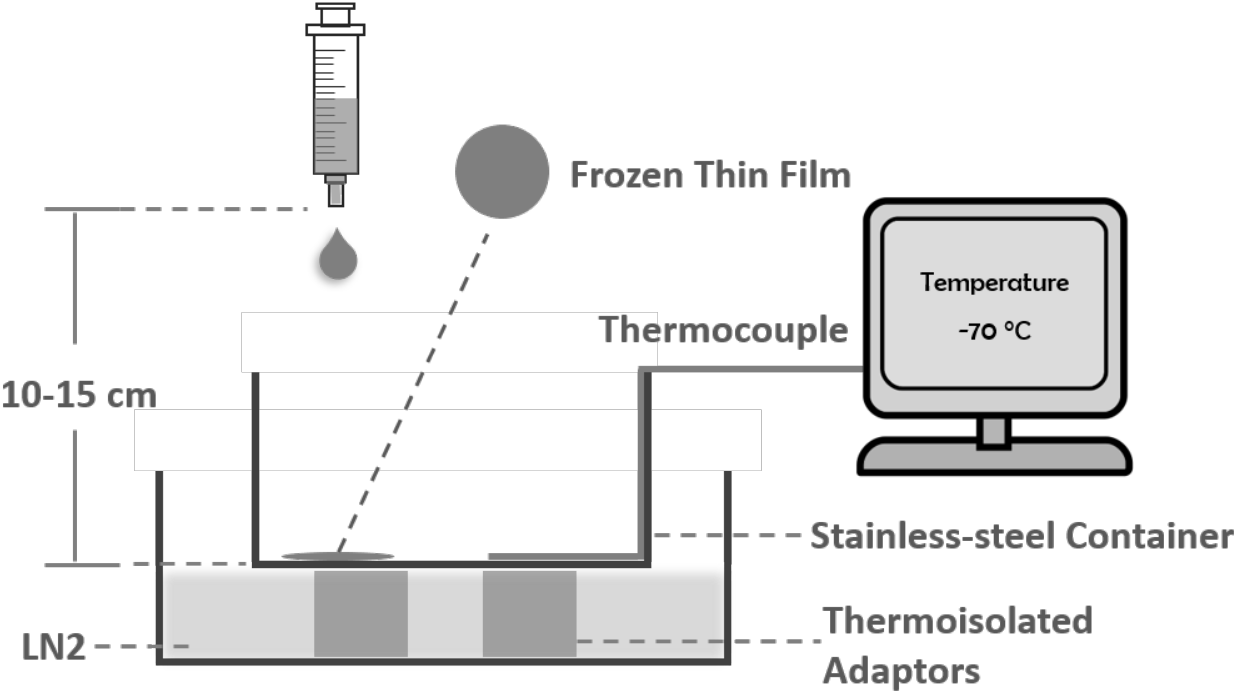
Graphical diagram of thin film freezing system. LN2: liquid nitrogen

### 5. TFFD Powder Characterization

#### 5.1 Particle size measurement

The geometric particle size distribution of the TFFD processed phage powders was analyzed using a Sympatec HELOS laser diffraction instrument (Sympatec GmbH, Germany) equipped with RODOS dry dispersion. Measurements were taken every 10 milliseconds following powder dispersion at 3 bar. Measurements that had an optical density between 5% and 25% were then averaged to determine the particle size distribution. The particle sizes by volume were reported at the 10^th^, 50^th^, and 90^th^ percentiles (i.e. Dv10/50/90, respectively), as well as the percentages of particles falling within the 1-5 µm size range. Span was calculated with following equation: Span = (Dv90-Dv10)/Dv50. The dry dispersion method was used to study how the TFFD powder compositions were able to overcome binding forces between the agglomerated particles contained in the brittle matrix powder, i.e. how easily the TFFD powder samples were able to be sheared into smaller particles of aggregates. This property is relevant for future DPI and nebulization studies with these powder compositions.

#### 5.2 Scanning electronic microscope (SEM)

The morphologies of the various TFFD processed phage powders were analyzed with Zeiss Supra 40VP SEM (Carl Zeiss Microscopy GmbH, Jena, Germany). Samples were mounted on aluminum SEM stubs using a carbon conductive tape and were coated with 15 nm of platinum/palladium (Pt/Pd) using a Cressington sputter coater 208 HR (Cressington Scientific Instruments Ltd., Watford, UK).

#### 5.3 X-ray diffraction (XRD) pattern

The crystallinity of TFFD processed phage powders was detected using an X-ray diffractometer (MiniFlex 600, Rigaku Co., Japan) under ambient conditions. Powders were spread on the glass slides and were exposed to Cu Kα radiation at 15 mA and 40 kV. The scattered intensity was collected by a detector in the 2θ range 5 to 50° with a step of 0.025° and a speed of 2°/min.

#### 5.4 Thermogravimetric analysis (TGA)

Thermogravimetric analysis was conducted using the Mettler Thermogravimetric Analyzer (Mettler Toledo, Columbus, OH, US). Samples weighing 1-3 mg were loaded in 70 µl alumina pans which were loosely capped with a lid containing a vent hole. Samples were heated up from 35 °C to 400 °C at a rate of 10 °C/min. The system was purged by nitrogen at a flow rate of 50 L/min. The percentage of change in mass over initial mass was calculated and plotted as a function of temperature. The percent of weight loss at 120 °C was used to identify the water content in powders. The reliability of TGA was confirmed by testing the same samples with an isothermal heating cycle, i.e. ramping from 35 °C to 120 °C at a rate of 10 °C/min followed by holding at 120 °C for 10 min. There was no significant difference between the two methods.

## Results and Discussion

### 1. Phage viability was affected by type and amount of excipient in the composition

#### 1.1 Formulation screening

The sugars that were tested, which included lactose, trehalose, and sucrose, were formulated with leucine at sugar: leucine weight ratios of 100: 0, 90:10, 75:25, and 60:40. The titer loss results are shown in Figure 1 where the preservation effect of buffers for T7 phage viability was demonstrated. In all tested groups, the titer losses from largest to smallest were the following: no buffer > PBS > SM, with only one exception where the titer in one SM containing formulation was slightly less than its PBS counterpart for sucrose: leucine 60: 40. All of the no buffer formulations exhibited a more than 2 log pfu titer reduction, while all of the buffer containing formulations, with the exception of lactose in the PBS formulation group (average of 2.23 log pfu titer reduction) exhibited less than 2 log pfu titer reduction. Moreover, there was less than 1 log pfu titer reduction in nearly all SM buffered formulations, with the exception of sucrose: leucine 60: 40, which exhibited an average titer reduction of 1.60 log pfu.

**Figure 1.**
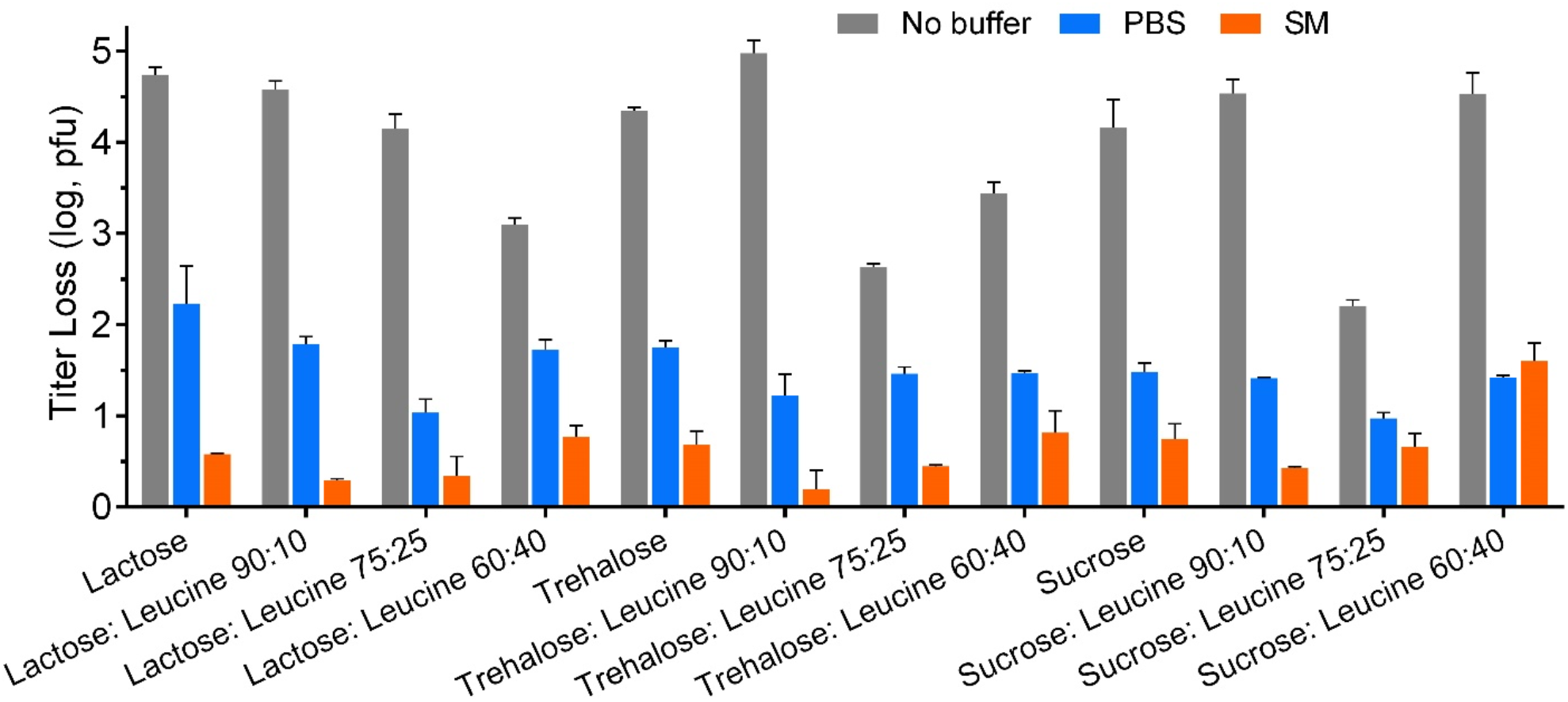
Titer loss of T7 phage after thin film freeze-dried in different formulations with or without buffer systems

Phage were most inactivated when trehalose was present in the composition, such as the trehalose: leucine 90:10 No buffer sample which averaged a 4.97 log pfu titer loss. In contrast, phage activity was best preserved in the trehalose: leucine 90: 10 SM buffered group, exhibiting the smallest titer loss, with an average of 0.19 log pfu titer reduction. Among all of the PBS samples, the sucrose: leucine 75:25 composition exhibited the best preservation of phage activity with an average titer loss of only 0.97 log pfu. In order to investigate the effect of the buffer systems on the TFFD phage formulations, the trehalose: leucine 90:10 and sucrose: leucine 75:25 groups were selected for further investigation.

#### 1.2 Thin film freeze drying processing preserves phage bioactivity

The TFFD process involves two major steps: ultra-rapid freezing and drying, both of which can be damaging to phage stability. Incorporating a buffer system into the TFFD compositions reduced the total titer loss in both freezing and drying steps regardless of excipient compositions. In order to investigate the ability of the excipient matrices to preserve phage activity, the viability of T7 phage after each processing step during the TFFD powder production was examined. The observed accumulated reduction in titers is shown in Figure 2. Generally, a greater titer loss was observed after the drying step than after the freezing step, which indicates that the stresses in the drying step were more detrimental to T7 phage. The differences of titer reduction between formulations were less pronounced after the ultra-freezing step and were greater after drying. The titer losses of T7 phage after both TFFD processing steps were greatly reduced by the presence of a buffer in the TFFD composition.

**Figure 2.**
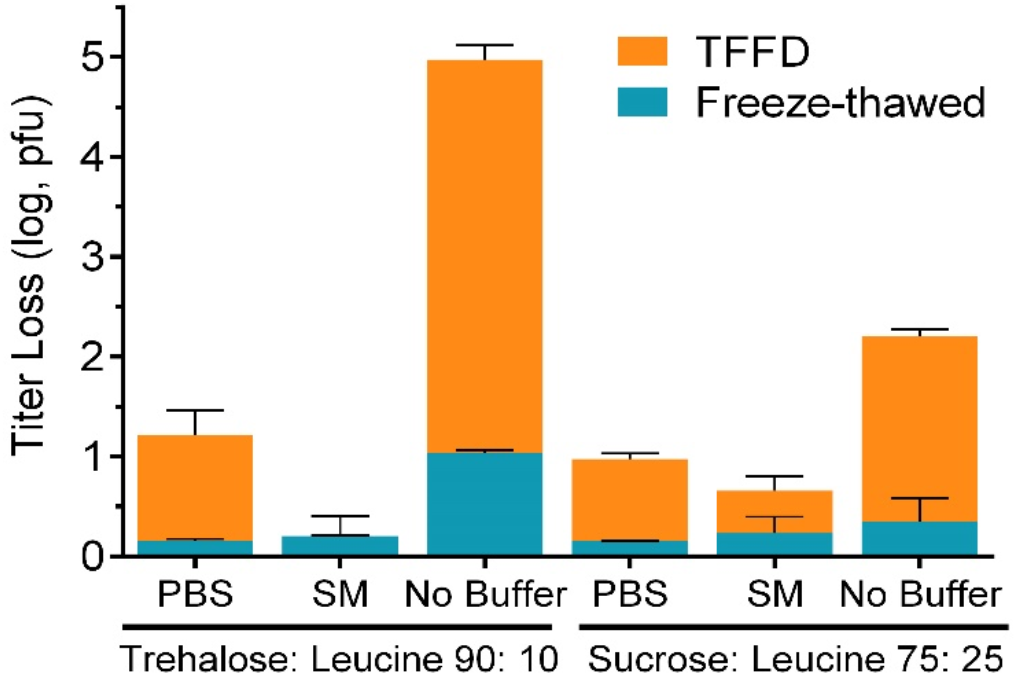
Titer loss of T7 phage after each processing step of thin film freeze-drying

Comparing the PBS and SM buffer systems, the cryoprotection (i.e., protection during the ultrarapid freezing step of the TFFD process) of the PBS containing compositions was slightly better than that of the SM containing compositions. The average titer losses after freezing were only 0.15-0.16 log pfu for the PBS containing formulations and 0.20-0.23 log pfu for the SM containing formulations. Interestingly, phage viability was preserved during the drying step when SM buffer was present in the TFFD composition. The formulation of trehalose: leucine 90:10 in SM buffer provided T7 phage with complete protection from dehydration stresses, as indicated by no titer loss (−0.01 log pfu titer loss) after the drying step. When combining the titer reductions accumulated from the two steps in the TFFD process, the average total titer losses of the buffer containing formulations were only 1.22 log pfu for the trehalose: leucine 90:10 PBS composition, 0.19 log pfu for the trehalose: leucine 90:10 SM composition, 0.97 log pfu for the sucrose leucine 75:25 PBS composition, and 0.66 log pfu for the sucrose: leucine 75:25 SM composition.

#### 1.3 Physical characterization of the TFFD powder compositions

##### 1.3.1 Geometric particle size distribution is consistent for TFFD powder compositions

The geometric particle size distributions of the TFFD phage powder compositions were determined by dispersing the dry powder followed by laser diffractometry measurements (Table 1). Each of the TFFD powder compositions were in the micron size range, with the average Dv50 less than 10 µm. In particular, the Dv50 was less than about 5 µm for the sucrose: leucine 75:25 SM powder and the PBS containing formulations. The percentage of particles falling within the 1-5 µm size range was greater than 40% in those powders. The particles were generally smaller in the sucrose: leucine 75:25 group than in the trehalose: leucine 90:10 group. One exception was the PBS containing formulations, in which the particles generated from the TFFD powders containing sucrose: leucine 75:25 were larger than in the trehalose: leucine 90:10 formulations. Compared to the powder compositions containing no buffer, the particle sizes were decreased in buffer containing powders in both excipient matrices and this size reduction was more significant in compositions containing the PBS buffer. However, the geometric particle size range of the PBS powders were greater than that of its no buffer and SM counterpart powder compositions. It is worth noting that typically a large geometric particle size (>5 µm) of TFFD powder is not necessarily indicative of a great aerodynamic particle size [29]. The particles generated by TFFD are brittle, can be easily dispersed with relatively low dispersion force into fine particles that falls into an inhalable particle size range (< 5 µm). Therefore, TFFD generated phage powders have great potential to be applied in respiratory delivery.

**Table 1.**
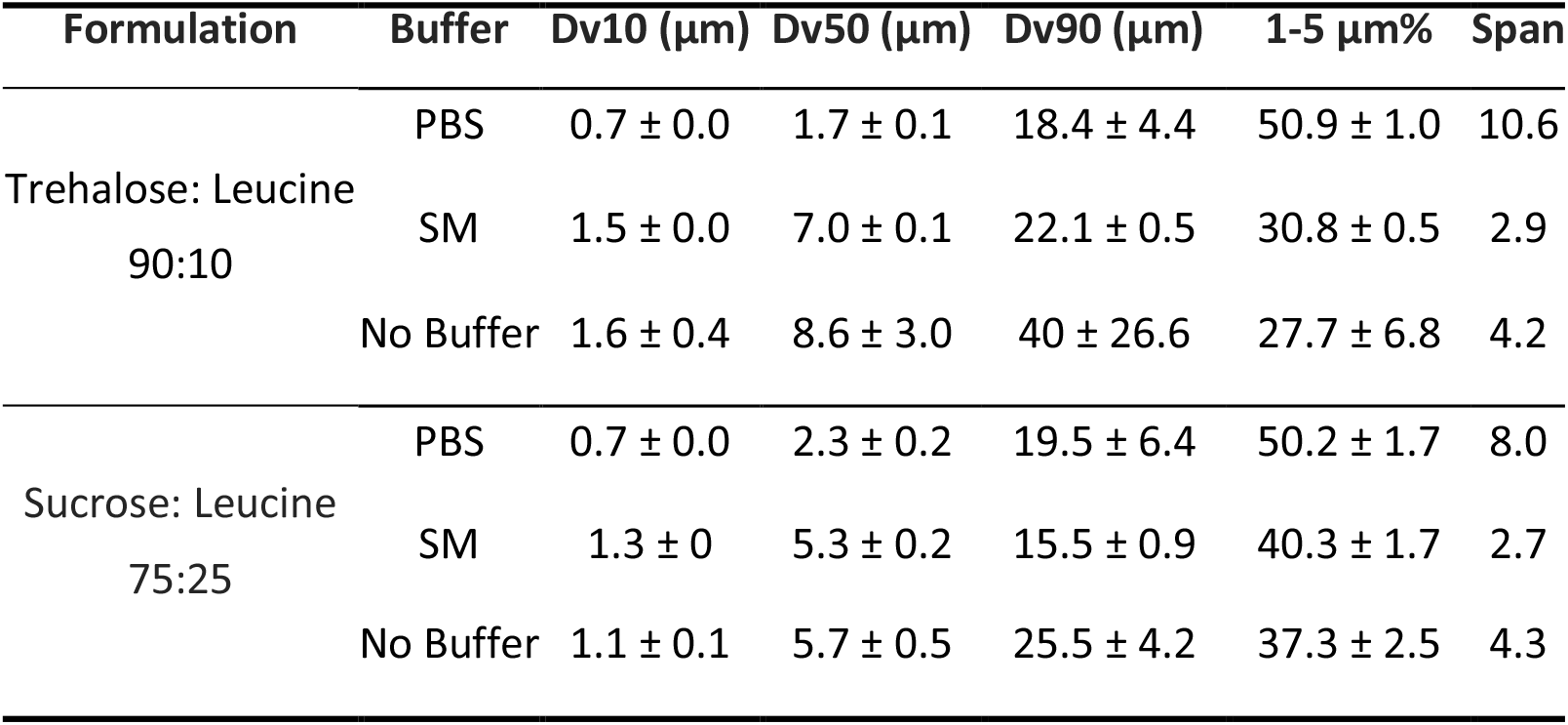
Particle size (distribution) of thin film freeze-dried T7 phage powders

| Formulation | Buffer | Dv10 ($\mu\text{m}$) | Dv50 ($\mu\text{m}$) | Dv90 ($\mu\text{m}$) | 1-5 $\mu\text{m}\%$ | Span |
| --- | --- | --- | --- | --- | --- | --- |
| Trehalose: Leucine<br>90:10 | PBS | $0.7 \pm 0.0$ | $1.7 \pm 0.1$ | $18.4 \pm 4.4$ | $50.9 \pm 1.0$ | 10.6 |
| | SM | $1.5 \pm 0.0$ | $7.0 \pm 0.1$ | $22.1 \pm 0.5$ | $30.8 \pm 0.5$ | 2.9 |
| | No Buffer | $1.6 \pm 0.4$ | $8.6 \pm 3.0$ | $40 \pm 26.6$ | $27.7 \pm 6.8$ | 4.2 |
| Sucrose: Leucine<br>75:25 | PBS | $0.7 \pm 0.0$ | $2.3 \pm 0.2$ | $19.5 \pm 6.4$ | $50.2 \pm 1.7$ | 8.0 |
| | SM | $1.3 \pm 0$ | $5.3 \pm 0.2$ | $15.5 \pm 0.9$ | $40.3 \pm 1.7$ | 2.7 |
| | No Buffer | $1.1 \pm 0.1$ | $5.7 \pm 0.5$ | $25.5 \pm 4.2$ | $37.3 \pm 2.5$ | 4.3 |

##### 1.3.2 Crystallinity of the TFFD is determined by the excipient composition

The crystallinity of the TFFD T7 phage powders was studied using XRD (Figure 3). All formulations exhibited a partially crystalline and amorphous structure, which depended on the type of excipient present in the composition. Characteristic peaks for NaCl were observed at 2θ degrees of 27.8°, 32° and 45.5° across all buffer containing formulations [30]. In addition to NaCl characteristic peaks, SM buffer samples also have some relatively shorter peaks at 2θ degrees of 10.9°, 15.7°, and 21.8° to 23.7°, 26°, 27.5°, 39.3° to 43°. These are likely due to other components used in the buffer, including Tris and MgSO_4_. The characteristic peaks for leucine, including peaks at 2θ degrees of 7°, 19.2°, and 24.6°, were more pronounced in the sucrose leucine 75:25 samples because it contained 15% more leucine as compared to the trehalose leucine 90:10 compositions [31]. No characteristic peaks were observed for sucrose and trehalose containing compositions, which indicate that these compositions were amorphous after the TFFD process. The TFFD powder compositions that did not contain buffer were amorphous as indicated by the broad ‘halo’ peak.

**Figure 3.**
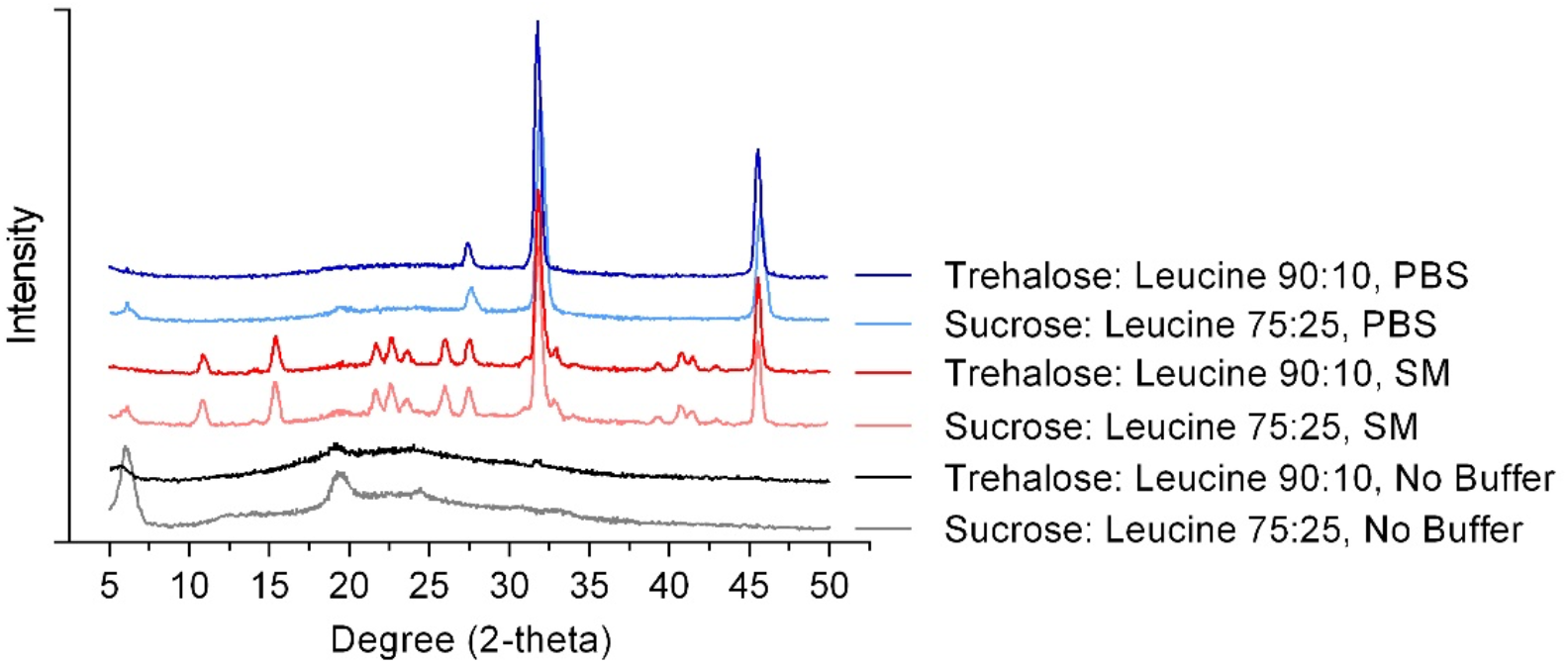
X-ray diffraction patterns of TFFD T7 phage powders

##### 1.3.3 Morphology of the TFFD powder compositions

TFFD phage powder morphologies were studied by SEM (See Figure 4). For all samples, SEM images indicated characteristic highly porous structures made up of submicron structured primary particles all interconnected within the matrix. The PBS and no buffer containing formulations exhibited a higher porosity as compared to the SM containing formulations, which was also reflected in the geometric particle size measurements described above. The sucrose: leucine 75:25 formulations resulted in smaller sized aggregates as compared to the trehalose: leucine 90:10 formulations.

**Figure 4.**
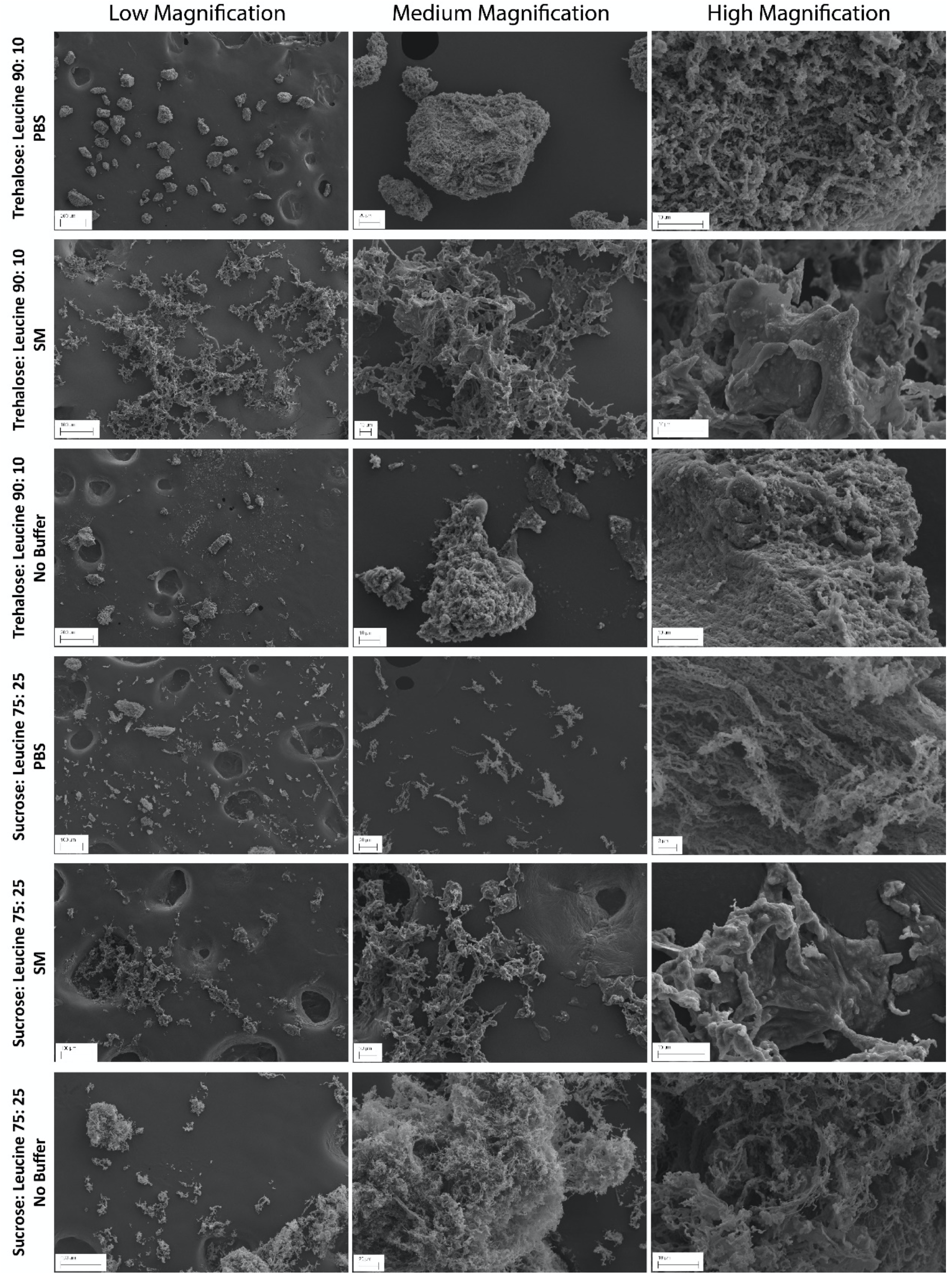
Scanning electron microscopy images of thin film freeze-dried T7 phage powders

Visually, the SM formulations exhibited powder characteristics different from those of the PBS and no buffer formulations, exhibiting a more continuous aggregated particle network when observed at low and medium magnification. At a lower magnification, SM formulations exhibited more thin and elongated shapes. At high magnification, SM formulations exhibited a noticeably smoother morphology with irregularly large shaped perturbances on the surface. Notably, the sucrose: leucine 75:25 SM formulations exhibited less particle networking as compared to the structures observed in the trehalose: leucine 90:10 SM formulations.

Lastly, for the sucrose: leucine 75:25, the formulations containing no buffer exhibited more fibrous networking at medium and high magnifications, while the trehalose: leucine 90:10 formulations appeared visually to be composed of denser, sponge-like structures with irregular flaky nano-structures attached onto their surfaces.

##### 1.3.4 Water content and thermal analysis of the TFFD powders

TGA was used to investigate the thermal stability profiles (Figure 5) and to determine the water content in the TFFD T7 phage powders (Figure 6). Most of the powders showed more than one mass loss steps since there were multiple components that underwent sublimation during heating, such as the sugars, leucine, and Tris (contained in the SM buffer). The onset of decomposition was at 170-180 °C and 205-215 °C for the sucrose: leucine 75:25 formulations and 205-215 °C and 270-280 °C for the trehalose: leucine 90:10 formulations, respectively. The TGA curves of the SM buffer containing formulations were different from their PBS and no buffer containing formulation counterparts.

**Figure 5.**
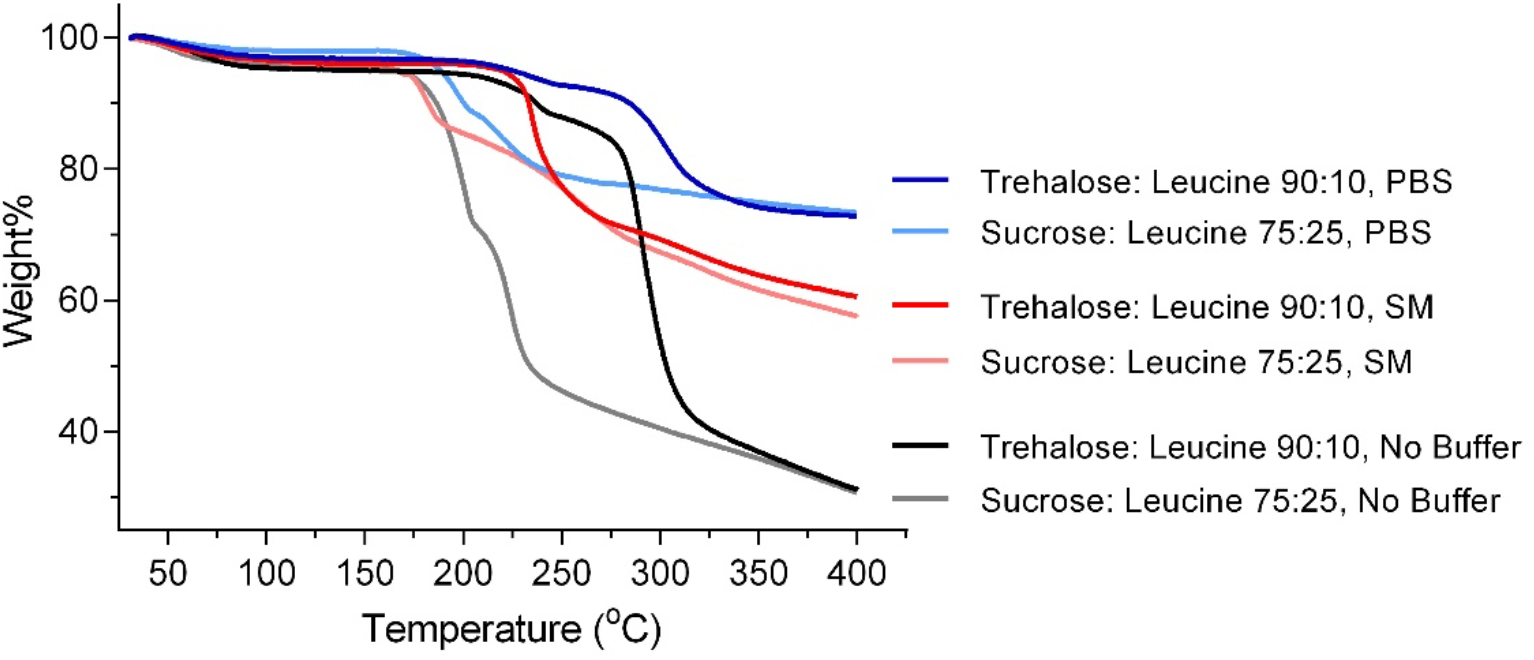
Thermogravimetric analysis curves of thin film freeze-dried T7 phage powders

**Figure 6.**
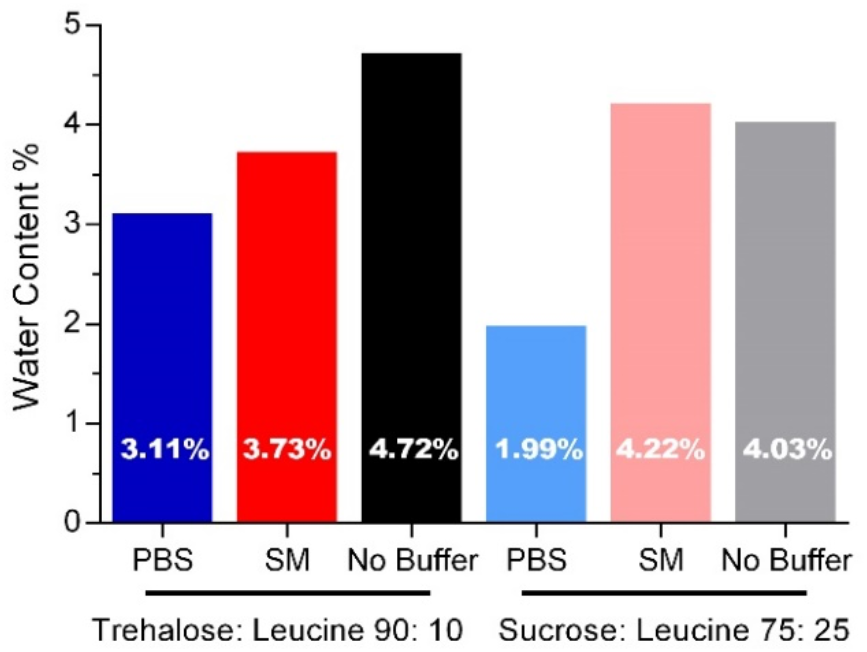
Water contents in thin film freeze-dried T7 phage powders

The water content was less than 5% in all TFFD powders and was generally lower in buffered formulations (Figure 6). PBS containing formulations resulted in the lowest water content. In comparison to formulations without buffer, the addition of PBS buffer to the TFFD compositions decreased the water content to 1.61% and 2.04% in the trehalose: leucine 90:10 and the sucrose: leucine 75: 25 formulations, respectively. Meanwhile, the effect of SM buffer on residual moisture content after TFFD processing differed between the two excipient matrices. The presence of SM buffer reduced the water content in the trehalose: leucine 90:10 formulation by about 1% while the water content was slightly higher for the sucrose: leucine 75:25 formulation.

## Conclusion

Thin film freeze drying was successfully used to prepare stable, readily shear-able brittle matrix powder phage formulations that have potential for use with various drug delivery systems, including dry powder inhalation and nebulization. Incorporation of a buffer system greatly enhanced the preservation of phage bioactivity, minimizing or eliminating any titer loss post TFFD-processing. Here we demonstrate that the thin film freezing process is a desirable particle engineering method to produce phage powder compositions because it eliminates the exposure of phage formulations to mechanical or thermal stresses that are encountered during spray drying, spray freeze drying and atmospheric spray freeze drying.

## Acknowledgements

Zhang and Williams acknowledge TFF Pharmaceuticals, Inc. for the financial support through a sponsored research agreement. Ghosh and Soto were supported by NIH National Heart, Lung and Blood Institute of the National Institutes of Health under award number R01HL138251. All authors are co-inventors on related intellectual property. The Board of Regents of The University of Texas has licensed IP related to research reported in this paper to TFF Pharmaceuticals, Inc. Williams acknowledges ownership of stock in TFF Pharmaceuticals, Inc.

